# 12-Lipoxygenase (12-LOX) Plays a Key Role in Hyperinflammatory Response Caused by SARS-CoV-2

**DOI:** 10.1101/2025.08.27.672781

**Authors:** Melinee D’silva, Matthew T. Vaughan, David J. Maloney, Sachin A. Gupte, Jerry L. Nadler, Chandra Shekhar Bakshi

## Abstract

The COVID-19 pandemic, caused by the SARS-CoV-2 virus, has led to significant global morbidity and mortality. The severe disease outcomes are often linked to a hyperinflammatory response known as a “cytokine storm.” The underlying mechanisms responsible for this exaggerated immune response remain incompletely understood. This study aimed to investigate the molecular pathways contributing to the severe inflammatory damage and mortality associated with COVID-19. SARS-CoV-2 hijacks host lipid metabolism, particularly the phospholipase A2 (PLA2) pathway, which leads to the production of the bioactive molecules, including the 12-Lipoxygenase (12-LOX)-derived lipid mediators in platelets, as well as in lung and vascular cells. We hypothesized that 12-LOX drives the hyperinflammatory response and disease severity, and that its inhibition could reduce inflammation and improve outcomes. Analysis of autopsy lung samples from COVID-19 decedents and SARS-CoV-2-infected K18-hACE2 transgenic mice revealed increased 12-LOX expression. We evaluated VLX-1005, a selective small-molecule 12-LOX inhibitor, in infected mice. Treatment initiated 48 hours post-infection significantly improved survival, reduced body weight loss, and decreased lung inflammation compared to controls. Notably, male mice showed higher survival rates than females. VLX-1005 treatment also suppressed key chemokines and cytokines associated with the cytokine storm, and reduced lung damage. These findings identify 12-LOX as a critical mediator of the hyperinflammatory response in severe COVID-19 and support its inhibition as a promising therapeutic strategy to mitigate inflammatory damage and reduce mortality.

**Significance:** This study provides critical insights into the mechanisms underlying severe COVID-19, identifying 12-Lipoxygenase (12-LOX) as a key driver of the hyperinflammatory response that contributes to disease severity and mortality. By demonstrating that SARS-CoV-2 hijacks host lipid metabolism to elevate pro-inflammatory lipid mediators, the research uncovers a novel pathogenic pathway that exacerbates lung inflammation. The use of VLX-1005, a selective 12-LOX inhibitor, significantly improved survival and reduced inflammatory damage in a mouse model, highlighting its therapeutic potential. These findings not only deepen our understanding of COVID-19 pathogenesis but also position 12-LOX as a promising target for intervention, offering a new avenue for mitigating the effects of cytokine storms in severe cases.

## Introduction

COVID-19, caused by Severe Acute Respiratory Syndrome Coronavirus 2 (SARS-CoV-2), has led to a global health crisis marked by high morbidity and mortality. While most infected individuals experience mild to moderate symptoms, a subset of patients develop severe disease characterized by acute respiratory distress syndrome (ARDS), multi-organ dysfunction, and, in some cases, death (1). A defining feature of severe COVID-19 is the development of a hyperinflammatory state known as a cytokine storm, which plays a central role in disease progression and tissue damage. The cytokine storm is an aberrant immune response characterized by the excessive and uncontrolled release of a plethora of immune modulators, including pro-inflammatory cytokines and chemokines. This hypercytokinemia leads to widespread immune cell infiltration, vascular leakage, and tissue injury, particularly in the lungs (2–4).

The lipoxygenase (LOX) family regulates inflammation through bioactive lipid mediators, with 12-LOX and 15-LOX (12/15-LOX in mice) playing distinct roles. Phospholipase A2 releases arachidonic acid, which reacts with 12-LOX to generate highly inflammatory lipids such as 12-S-hydroxyeicosatetraenoic acid (12-HETE) (5). 12-LOX, also known as “platelet type,” is encoded by *ALOX12*/*Alox12*, is expressed in platelets, epithelial cells, and macrophages. The enzymatic activity of the 12-LOX catalyzes the oxygenation of the 12^th^ carbon of the arachidonic acid, yielding the lipid molecule 12-HETE. In contrast, 15-LOX (*ALOX15*/*Alox15*), also known as “leukocyte type,” generates both 12-HETE and 15-HETE and is expressed in immune cells, neurons, and epithelial tissues (6). Intracellular production of 12-HETE induces oxidative stress, whereas extracellular 12-HETE modulates various signaling and inflammatory pathways (7). 12-LOX expression is elevated in obesity and metabolic syndrome. The 12-LOX-mediated detrimental inflammatory effects of obesity are linked to type 2 diabetes (T2D) and inflammation (8). Additionally, 12-LOX and 12-HETE have been associated with the development of Type 1 diabetes and T2D (9), hepatic and gastrointestinal inflammation, cardiovascular disease, neuroinflammation, and neurodegenerative diseases (10). Platelet activation, a critical process in thrombotic disorders like myocardial infarction and stroke, is also enhanced by 12-LOX activation, and selective inhibition of 12-LOX has been shown to reduce thrombosis (11). Activation of platelets by 12-LOX also leads to excessive clotting (12).

12-LOX plays a pivotal role in pulmonary inflammation, increased neutrophil influx, and cytokine production. Studies have demonstrated that 12-LOX induces an uncontrolled influx of neutrophils in the lungs of mice infected with *Streptococcus pneumoniae* (13) and triggers acute lung injury following inhalation of lipopolysaccharide (14). Intranasal administration of 12-LOX in mice leads to airway epithelial injury and airway hyperresponsiveness (15). Conversely, depletion of 12-LOX reduces lung inflammation by diminishing the recruitment of eosinophils, lymphocytes, and macrophages and lowering cytokine production and mucus secretion (16). Additionally, 12-LOX has been associated with asthma in children (17). In tuberculosis (18) and *Aspergillus fumigatus* (19) infections, increased expression of 12-LOX is associated with an increased neutrophil count and bacterial and fungal load, respectively, in the lungs.

The precise role of 12-LOX in the pathogenesis of COVID-19 remains unclear. However, significantly elevated levels of 12-HETE in the bronchoalveolar lavage fluid of COVID-19 patients compared to healthy controls suggest its involvement in the hyperinflammatory response observed in the lungs (20, 21). Studies have indicated that an increase in the levels of polyunsaturated fatty acids correlates with the severity of COVID-19 (21). A lipidomic storm with altered lipids, including 12-HETE, correlates with the inflammatory response and severity of COVID-19 (22). These findings suggest that the 12-LOX pathway may function upstream of the cytokine storm, which precedes the development of ARDS, respiratory failure, and death. The interplay between COVID-19 and diabetes has emerged as a significant clinical concern, with evidence supporting a bidirectional relationship. Individuals with preexisting diabetes are at heightened risk for severe COVID-19 outcomes, including hospitalization and mortality, due to underlying immune and metabolic dysregulation. Conversely, SARS-CoV-2 infection has been associated with an increased incidence of new-onset diabetes, likely driven by systemic inflammation, direct pancreatic injury, and treatment-related factors such as corticosteroid use (23). The objectives of this study were to determine whether SARS-CoV-2 infection upregulates 12-lipoxygenase (12-LOX) activity, thereby contributing to hyperinflammation, tissue injury, and mortality, and to evaluate the therapeutic efficacy of VLX-1005, a selective 12-LOX inhibitor, in attenuating inflammation and improving disease outcomes in a preclinical model of SARS-CoV-2 infection.

## Materials and Methods

### Human tissue samples

Tissue sections from lung and pancreas autopsy samples of deceased COVID-19 patients were obtained from the Columbia University BioBank. According to the New York Medical College (NYMC) Institutional Review Board policies, the use of these de-identified, processed human tissue sections is classified as “not human subjects research.” This classification is based on the fact that the study did not involve any intervention or interaction with the individuals, nor did it include any identifiable patient information.

### Mouse experiments

The K18-hACE2 transgenic mouse model, which recapitulates both mild and severe forms of human COVID-19 (4), was used in this study. Ten-week-old male and female K18-hACE2 mice (B6.Cg-Tg(K18-ACE2)2Prlmn/J, Stock No: 034860; The Jackson Laboratory, Bar Harbor, ME, USA) were intranasally infected with 1 × 10³, 2 × 10³, or 1 × 10⁵ PFU of SARS-CoV-2 (2019-nCoV/USA-WA1/2020; BEI Resources, Manassas, VA). Mice were anesthetized with intraperitoneal ketamine/xylazine, and the virus was administered intranasally (10 μL per nare). Immediately post-infection, mice were divided into two treatment groups: Group I received vehicle control (5% DMSO), and Group II received VLX-1005 (30 mg·kg⁻¹·day⁻¹), a specific inhibitor of 12-LOX (Kindly provided by Veralox Therapeutics Inc., Frederick, MD) via intraperitoneal injection for 7 days. In a separate experiment, treatment with vehicle or VLX-1005 was initiated on day 2 post-infection and continued through day 14. Mice were monitored daily for morbidity (body weight) and mortality. Humane endpoints were applied for mice exhibiting >20% body weight loss or signs of severe illness. Lung tissues were collected from mice that succumbed during the observation period or were euthanized at the study endpoint. Samples were preserved in Trizol for RNA isolation or in 10% formalin for immunofluorescence staining and histopathological analysis. All procedures were conducted in the Biosafety Level 3 laboratory at the Rutgers Regional Biocontainment Laboratory, New Jersey Medical School. All animal protocols were approved by the Rutgers Institutional Animal Care and Use Committee and all experiments were conducted according to NIH and institutional guidelines.

### Immunofluorescence staining

Immunofluorescence staining for phosphorylated 12-LOX and 15-LOX was performed on 5-μm paraffin-embedded lung tissue sections from both human samples and experimental mice. Sections were baked at 60°C for 1 hour, followed by deparaffinization in 100% xylene (3 × 4 min), 100% ethanol (3 × 2 min), 95% ethanol (2 × 2 min), and 70% ethanol (1 × 2 min), then rinsed twice in distilled water on a shaker. Heat-induced antigen retrieval was carried out by microwaving slides in citrate buffer (10 mM citrate, 0.05% Tween 20, pH 6.0). After cooling to room temperature, slides were rinsed in distilled water and blocked with 2% donkey serum for 30 minutes. Primary antibody incubation was performed overnight at 4°C using anti-12-LOX and 15-LOX antibodies for human tissues, and Rabbit anti-12-LOX and 15-LOX antibodies (MyBioSource) for mouse tissues. The next day, slides were washed in PBS (3 × 5 min) and incubated with Alexa Fluor 594-conjugated goat anti-rabbit IgG (A11005, 1:500 dilution) for 2 hours at room temperature. Nuclei were counterstained with DAPI. Images were acquired using a Zeiss LSM 980 confocal microscope with Airyscan II. Sections stained with secondary antibody alone served as negative controls.

### NanoString analysis

Mouse (n = 5 per group) and human (n = 5 per group) lung tissue were fixed in 10% formalin and submitted to the core services at the Albert Einstein College of Medicine for processing. Total RNA was extracted using standard protocols, and RNA concentrations were quantified using the Qubit fluorometric system. Gene expression analysis was performed using the NanoString nCounter platform (www.nanostring.com) with a custom Host Response Panel Code Set, profiling 785 genes across over 50 pathways in mouse and human samples. RNA expression was quantified using the nCounter Digital Analyzer, and both raw and normalized counts were generated using the Rosalind platform (https://app.rosalind.bio) and NanoString’s nSolver analysis software.

### Histopathological analysis

The superior and inferior lobes of the left lung from the experimental mice were fixed in 10% formalin. Following fixation, tissues were processed, paraffin-embedded, and sectioned at five μM thickness. Sections were then stained with hematoxylin and eosin (H&E) for histopathological evaluation in a blinded fashion.

### Statistical analysis

Statistical analyses were performed using GraphPad Prism version 9. Data are presented as mean ± standard error of the mean (SEM). Comparisons between two groups were conducted using unpaired two-tailed Student’s *t*-tests. Multiple groups were compared using two-way ANOVA, followed by Sidak’s post hoc multiple comparisons test. Post hoc analyses were performed only when the ANOVA indicated a statistically significant difference (*P* < 0.05). A *P* value < 0.05 was considered statistically significant.

## Results

### SARS-CoV-2 infection increases ALOX12 and immune gene expression in lung tissues from diabetic individuals

Immunofluorescence staining of lung tissue sections from diabetic individuals who succumbed to COVID-19 or unrelated respiratory illnesses was performed to assess 12-LOX and 15-LOX expression. The analysis revealed markedly elevated 12-LOX protein levels in COVID-19-positive patients compared to COVID-19-negative diabetic controls. Strong staining was observed in both alveolar and bronchial epithelial cells, whereas lungs from COVID-19-negative diabetic individuals exhibited only minimal 12-LOX staining (**Fig. 1A**). In contrast, no enhanced staining for 15-LOX was detected in COVID-19-positive diabetic lung tissue compared to COVID-19-negative samples (**Fig. 1B**).To complement the protein-level findings, gene expression profiling was conducted using RNA extracted from formalin-fixed, paraffin-embedded (FFPE) autopsy lung samples. Analysis via the NanoString nCounter platform revealed significantly increased *ALOX12* transcript levels in COVID-19-positive diabetic lungs relative to COVID-19-negative diabetic controls (**Fig. 1C**), indicating transcriptional activation of the 12-LOX pathway in response to SARS-CoV-2 infection. In addition to *ALOX12*, COVID-19 infection induced robust upregulation of several inflammatory and immune response genes. Transcripts for chemokine receptors *CXCR2, CCR7,* and *IL6R* were significantly elevated in COVID-19-positive diabetic lung tissue. Furthermore, interferon-stimulated genes *IFIT3* and *OASL* were significantly upregulated, with *STAT2* showing moderate elevation (Fig. 1D). These transcriptional changes suggest that the enhanced expression of *ALOX12* is driven by SARS-CoV-2 infection rather than diabetes alone. The concurrent upregulation of *ALOX12* and immune-related genes points to a potential role for 12-LOX in amplifying the inflammatory response in diabetic lungs during COVID-19. To determine whether 12-LOX upregulation occurs independently of diabetes, lung sections from deceased non-diabetic individuals with COVID-19 were also stained for 12-LOX. Positive staining confirmed that SARS-CoV-2 infection alone is sufficient to induce elevated 12-LOX expression (**Fig. 1E**). Collectively, these findings indicate that SARS-CoV-2 infection exacerbates 12-LOX expression in the lungs.

**Fig 1:**
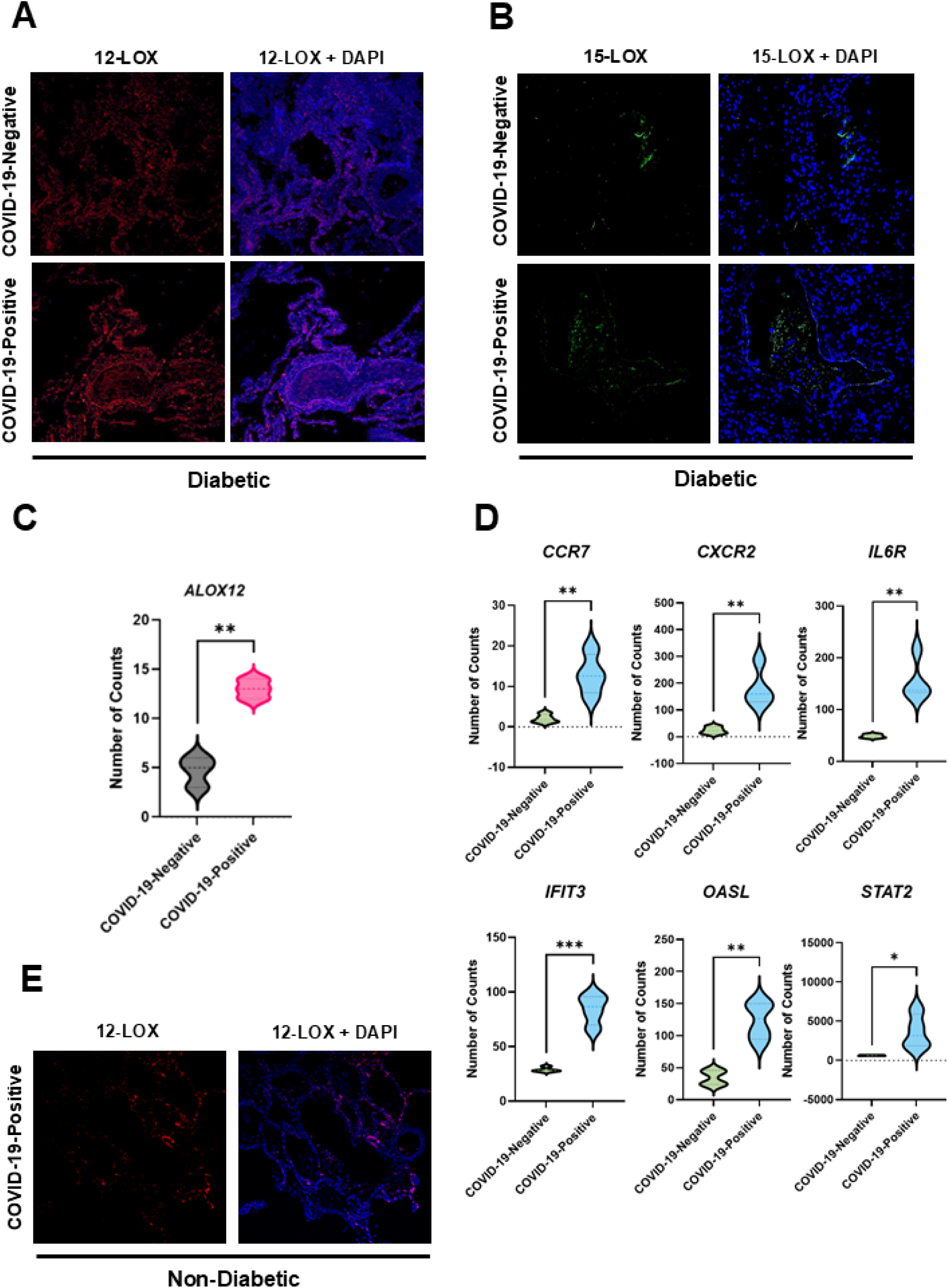
Increased pulmonary 12-LOX expression and inflammatory gene induction in COVID-19-positive diabetic lung tissue. **(A)** 12-LOX immunofluorescence staining of lung sections from diabetic COVID-19-negative and COVID-19-positive decedents. Images depict one of five tissue samples per group. 12-LOX is visualized in red, and nuclear counterstaining with DAPI is merged in purple (Magnification × 40×). **(B)** 15-LOX immunofluorescence staining of lung sections from diabetic COVID-19-negative and COVID-19-positive decedents. Images depict one of five tissue samples per group. 15-LOX is visualized in green, and nuclear counterstaining with DAPI (Magnification × 40×) **(C, D)** Gene expression analysis of inflammation-associated transcripts in lung tissue was performed using NanoString nCounter. **(C)** Quantitative *ALOX12* mRNA counts from deceased COVID-19-negative diabetic and COVID-19-positive diabetic patients (n = 5 per group). **(D)** Relative mRNA expression of the indicated genes in deceased COVID-19-negative diabetic and COVID-19-positive diabetic patients (n=5 per group). **(E)** 12-LOX immunofluorescence staining of lung sections from non-diabetic COVID-19-positive decedents. 12-LOX is visualized in red, and nuclear counterstaining with DAPI is merged in purple (Magnification × 40×). Statistical significance was determined by unpaired t-test (*p < 0.005; **p < 0.01; ***p < 0.001).

### SARS-CoV-2 infection increases 12-LOX expression in the lungs of non-diabetic K18-hACE2 mice

To determine whether SARS-CoV-2 infection alone is sufficient to induce pulmonary 12-LOX expression independent of metabolic disease, lung tissues from K18-hACE2 transgenic mice were analyzed following intranasal inoculation with a lethal viral dose (1 × 10⁵ PFU). Animals were necropsied immediately after death, and formalin-fixed lung sections were subjected to immunofluorescence staining for 12-LOX. In uninfected control mice, the 12-LOX signal was minimal. In contrast, SARS-CoV-2-infected mice demonstrated markedly enhanced 12-LOX staining distributed throughout alveolar and bronchial epithelial compartments (**Fig. 2A**). Notably, a slight increase in 15-LOX staining was observed in SARS-CoV-2-infected mice; however, the intensity of staining was considerably lower than that observed for 12-LOX (**Fig. 2B**). These observations confirmed that SARS-CoV-2 infection directly induces 12-LOX expression in the lung, independent of diabetes. When considered alongside the transcriptomic data from diabetic human lung tissues shown in Figure 1, these findings support the notion that SARS-CoV-2 infection triggers 12-LOX activity as part of the pulmonary inflammatory response.

**Fig 2:**
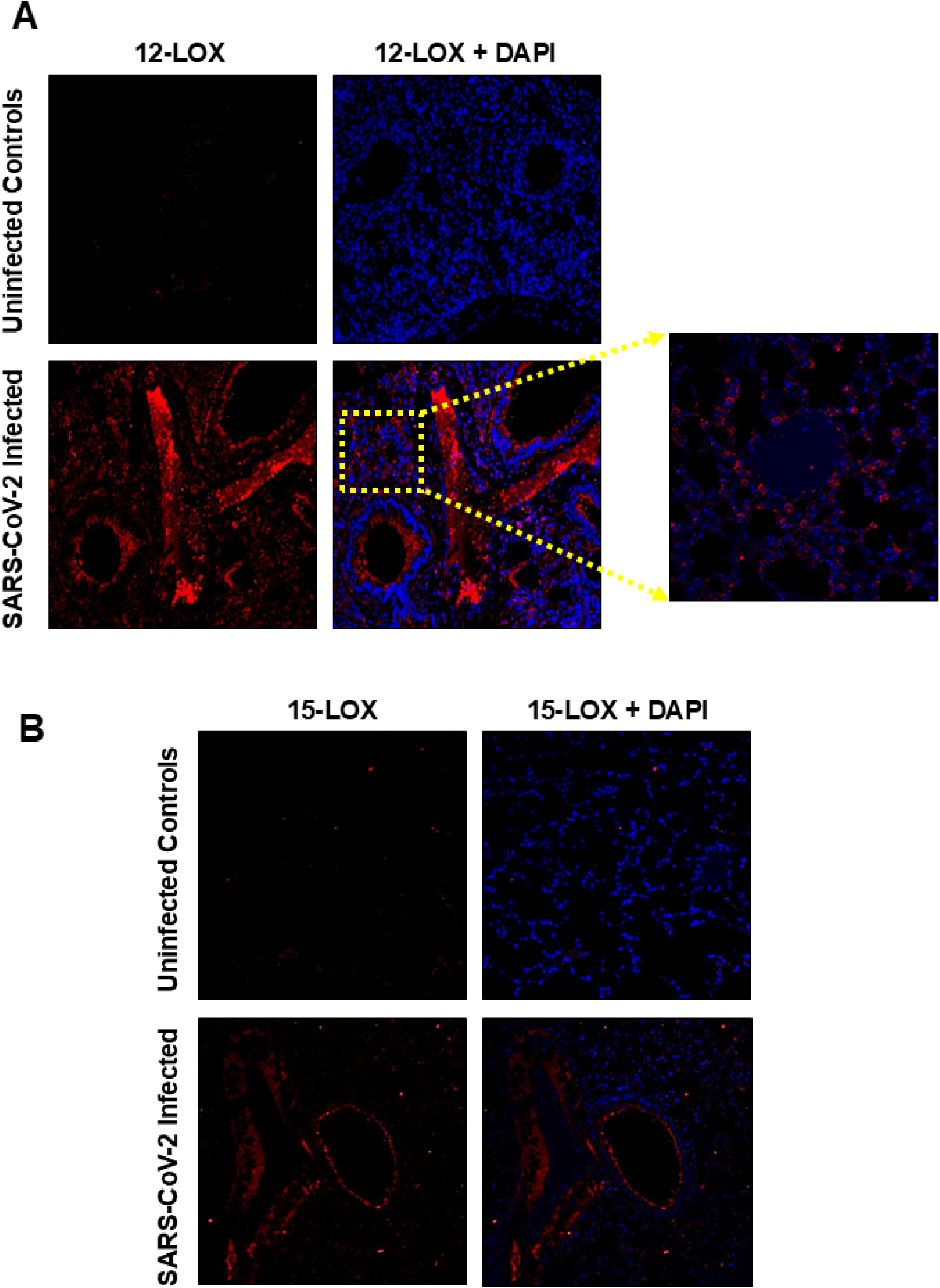
Increased pulmonary 12-LOX expression in SARS-CoV-2 K18-hACE2 mice. Immunofluorescence staining of lung sections for (**A**) 12-LOX and (**B**) 15-LOX expression in K-18hACE2 mice infected intranasally with 1×10^5^ PFUs of SARS-CoV-2. The mice were necropsied immediately after death, and lung tissues were fixed in 10% formalin, paraffin-embedded, and sectioned. Images depict one of five tissue samples per group. (Magnification × 40×; Inset 60×).

### Inhibition of 12-LOX activity reduces SARS-CoV-2-induced morbidity and mortality in K18-hACE2 mice

Based on the results shown in Figures 1 and 2, we hypothesized that increased 12-LOX activity contributes to the severity of SARS-CoV-2 infection and its inhibition would mitigate the severity of the infection. We evaluated whether pharmacologic inhibition using a specific 12-LOX inhibitor, VLX-1005, could reduce morbidity and mortality in SARS-CoV-2-infected mice. Male and female K18-hACE2 transgenic mice were infected intranasally with increasing doses of SARS-CoV-2 and treated with VLX-1005 via daily intraperitoneal injection during the indicated periods, as shown in Figure 3. Body weight and survival were monitored throughout the study period. No differences were observed between untreated and VLX-1005-treated male or female mice infected with 1 × 10³ PFU SARS-CoV-2 and treated with VLX-1005 beginning immediately after infection for seven consecutive days. An equal number of mice from both groups survived this infection dose with minimal morbidity, as indicated by their body weights (**Fig. 3A**). Next, we increased the infection dose. Mice were infected intranasally with 2 × 10³ PFU of SARS-CoV-2, and VLX-1005 treatment was initiated at 2 days post-infection and continued for 14 days. Female mice showed lower disease severity with only a slight improvement in survival following treatment, as evidenced by preserved body weight and 100% survival. However, male mice receiving VLX-1005 demonstrated improved outcomes, including significantly higher survival rates and reduced weight loss. Survival differences between VLX-1005-treated and vehicle control male mice were statistically significant (log-rank test, p < 0.05) (**Fig. 3B**). Since female mice showed reduced susceptibility to the 1 × 10³ and 2 × 10³ PFU infection doses of SARS-CoV-2, the viral dose was increased to 1 × 10⁵ PFU in a subsequent experiment. The therapeutic efficacy of VLX-1005 was determined by treating infected mice from day 2 to day 14 post-infection. Both vehicle control and the VLX-1005-treated groups exhibited rapid weight loss for 7-8 days post-infection. However, treatment with VLX-1005 improved survival in female mice. The VLX-1005-treated group regained their body weights, and 60% mice survived the higher infection dose (**Fig. 3C**). These findings demonstrate that the inhibition of 12-LOX using VLX-1005 reduces disease severity and improves survival of SARS-CoV-2-infected mice, indicating that its inhibition has protective effects.

**Fig 3.**
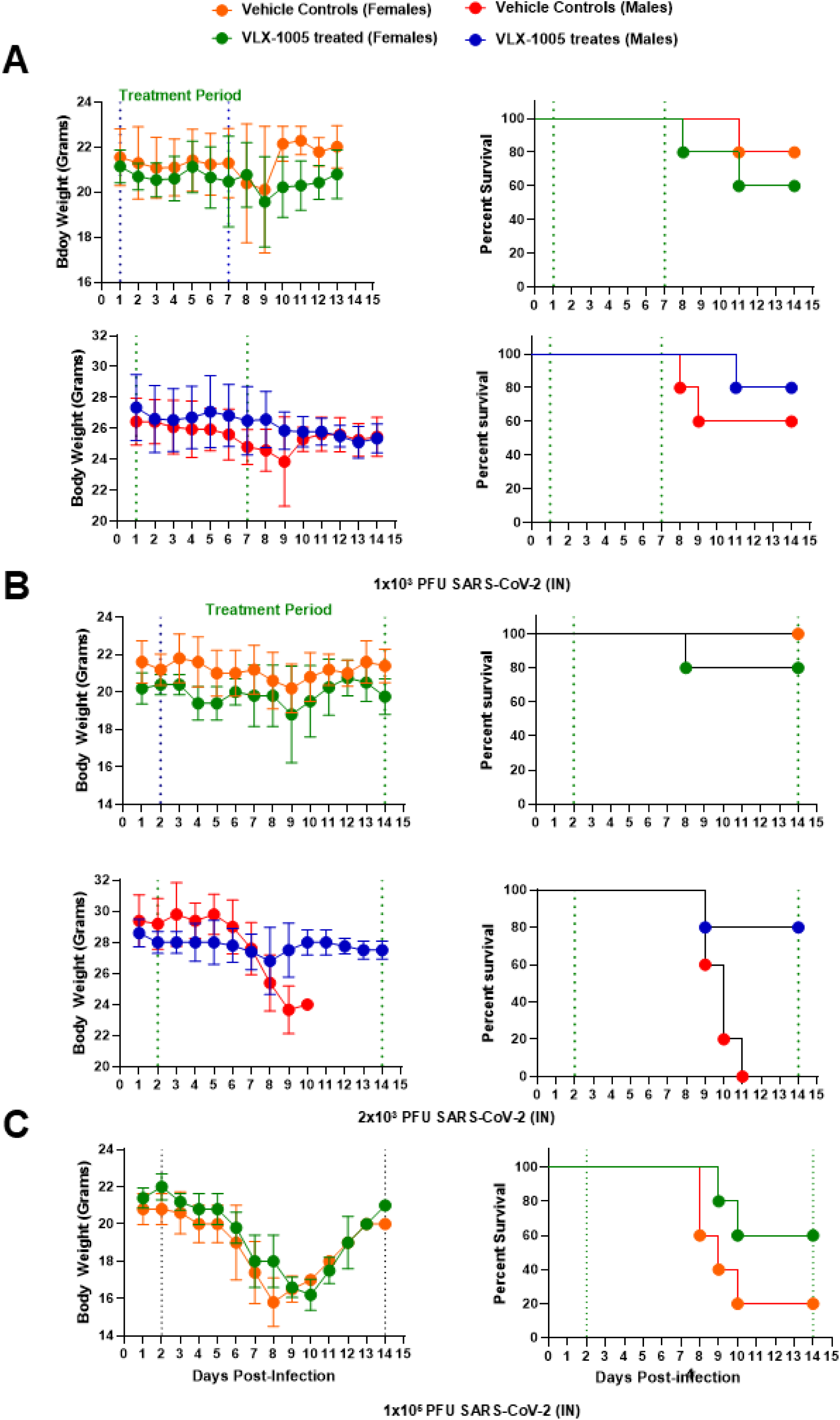
Inhibition of 12-LOX activity reduces SARS-CoV-2-induced morbidity and mortality in K18-hACE2 mice. Male and female K18-hACE2 transgenic mice (n = 10 mice per group) were intranasally (IN) inoculated with **(A)** 1 × 10³, **(B)** 2 × 10³, or **(C)** 1 × 10⁵ plaque-forming units (PFU) of SARS-CoV-2. Mice were treated with the 12-lipoxygenase (12-LOX) inhibitor VLX-1005 (30 mg/kg/day, intraperitoneally) either immediately post-infection for seven consecutive days **(A)** or starting two days post-infection and continuing through day 14 (**B and C**). Mice were monitored daily for morbidity, assessed via body weight measurements, and mortality. Survival data were analyzed using Kaplan-Meier survival curves, and statistical comparisons between groups were performed using the log-rank (Mantel-Cox) test. Vertical dotted lines indicate the treatment window.

### Inhibition of 12-LOX reduces inflammatory gene expression and lung pathology in SARS-CoV-2-infected male mice

To investigate the molecular impact of 12-LOX inhibition during SARS-CoV-2 infection, male K18-hACE2 mice were intranasally infected with 2 × 10³ PFU of SARS-CoV-2 and treated daily with either VLX-1005 or vehicle from day 2 to day 14 post-infection. Gene expression analysis using the NanoString nCounter platform revealed that VLX-1005 treatment significantly reduced *12lox* mRNA levels compared to vehicle-treated controls, indicating effective suppression of 12-LOX (**Fig. 4A**). To assess the enzymatic activity of 12-LOX, 12-HETE levels, a functional marker of 12-LOX, were quantified by ELISA. VLX-1005 treatment led to a marked reduction in 12-HETE concentrations, confirming inhibition of 12-LOX at both the transcriptional and functional levels (**Fig. 4B**). NanoString profiling further revealed differential expressions of immune and inflammatory genes. In VLX-1005-treated, SARS-CoV-2-infected mice, 30 genes were downregulated and four were upregulated relative to vehicle-treated controls (**Fig. 4C**). Notably downregulated genes included chemokines (*Ccl11, Ccr7, Ccrl2, Cd14, Cxcl2, Cxcl5,* and *Cxcr2*), cytokine-related genes (*Ifit1, Ifit3, Il1rap, Il1r2,* and *Il6ra*), and immune modulators (*Map3k8, Mefv, Ms4a4a, Rsad2, Socs3* and *Tap1*). Conversely, *Fos, Klrd1,* and *Tgfβ* were among the few genes upregulated following VLX-1005 treatment (**Fig. 4D**). Importantly, transcripts of *Nfkb1* and *Nfkb2*, key components of canonical and non-canonical NF-κB signaling pathways, were significantly reduced, suggesting diminished inflammatory transcriptional activity (**Fig. 4E**). Histopathological examination of lung tissues supported these molecular findings. While uninfected mice displayed normal lung architecture, vehicle-treated infected mice exhibited extensive alveolar damage, inflammatory infiltration, and structural disruption. In contrast, VLX-1005-treated mice showed preserved alveolar integrity and reduced inflammatory cell infiltration (**Fig. 4F**). Collectively, these findings demonstrate that VLX-1005 effectively suppresses 12-LOX expression and activity, attenuates pro-inflammatory gene expression, and mitigates lung pathology in SARS-CoV-2-infected mice. These results demonstrate the role of 12-LOX as a key driver of pulmonary inflammation and highlight its potential as a therapeutic target in COVID-19-associated lung injury.

**Fig 4.**
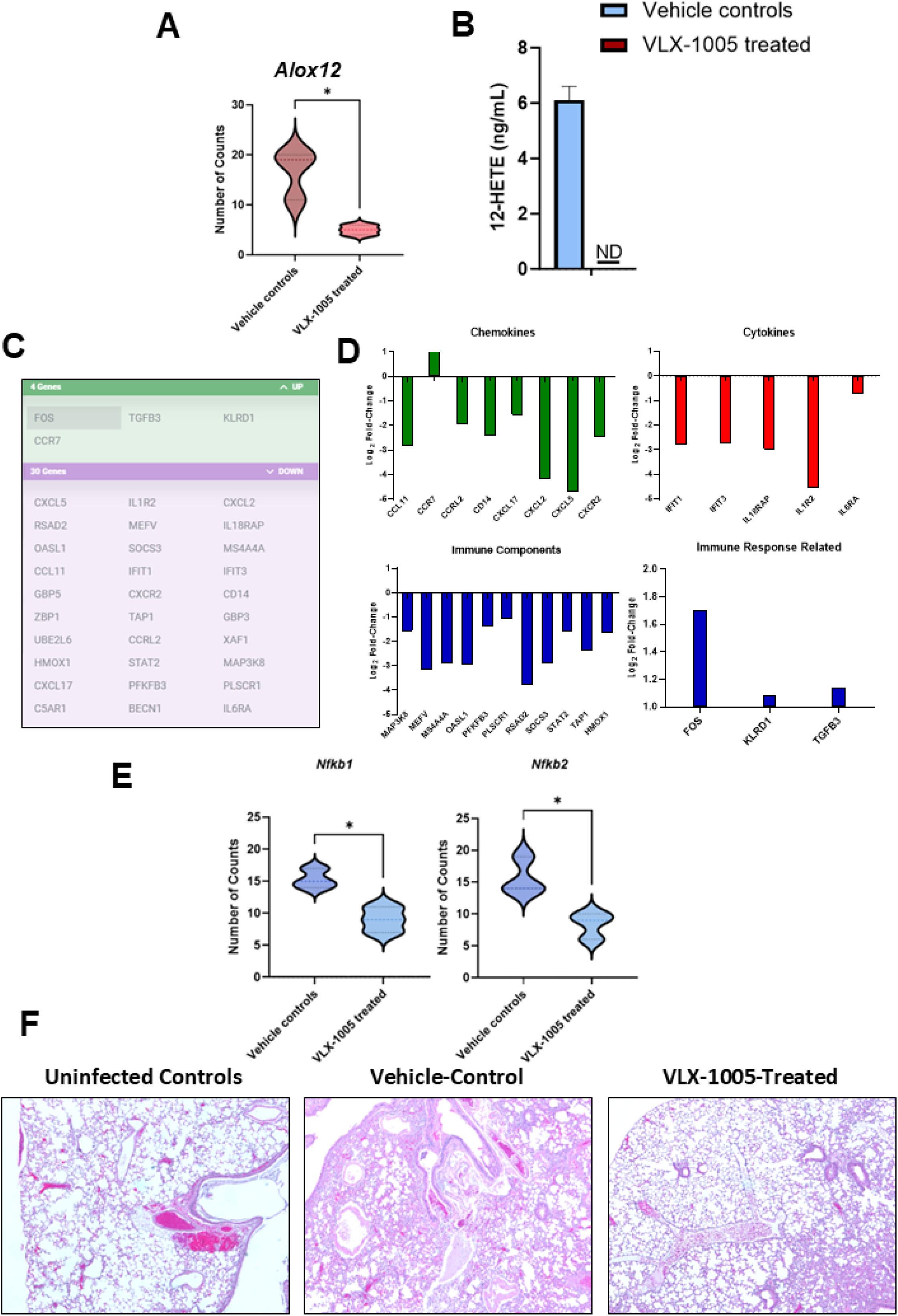
Inhibition of 12-LOX reduces inflammatory gene expression and lung pathology in SARS-CoV-2-infected male mice. Male K18-hACE2 transgenic mice were intranasally infected with 2 × 10³ PFU of SARS-CoV-2 and treated intraperitoneally with either vehicle or VLX-1005 (30 mg/kg/day) from 2 to 14 days post-infection. All VLX-1005-treated mice were euthanized, while vehicle-treated mice were necropsied immediately upon mortality for sample collection. Lung tissues and serum were harvested for molecular and histological analyses. **(A)** NanoString quantification of *Alox12* mRNA levels in lung tissues (n = 3 mice per group). **(B)** 12-HETE levels in serum as determined by ELISA (n = 5 mice per group). **(C)** Summary of differentially expressed genes identified by NanoString analysis (n = 3 mice per group). Log₂ fold-change in expression of indicated chemokines, cytokines, and immune response-related genes **(D)** and NF-κB1 and NF-κB2 in lungs **(E)** (n = 3 mice per group). The data were analyzed using Student’s *t-test.* *p<0.05; **p<0.01. **(F)** Representative H&E-stained lung sections from uninfected controls, SARS-CoV-2-infected vehicle-treated, and VLX-1005-treated mice (n = 3 mice per group).

### 12-LOX inhibition alters immune gene expression in female mice infected with SARS-CoV-2

To further evaluate the impact of 12-LOX inhibition, female K18-hACE2 mice were infected with a higher dose of SARS-CoV-2 (1 × 10⁵ PFU) and treated daily with VLX-1005 or vehicle from days 2 to 14 post-infection. Consistent with findings in male mice, VLX-1005 treatment significantly reduced *Alox12* gene expression and its lipid mediator 12-HETE in female mice, confirming effective inhibition of the 12-LOX pathway (**Fig. 5A, B**). However, transcriptional profiling revealed a distinct gene expression signature in female mice compared to males. Among the significantly downregulated genes were *Cxcl2, Cxcl1, Lyn, Plg, Plekhat, Prf1, Map2k3,* and *Lrrk2*. Interestingly, *Pfkfb3* was the only gene significantly upregulated following VLX-1005 treatment (**Fig. 5C**). Similar to male mice, transcripts of *Nfkb1* and *Nfkb2* were significantly reduced in VLX-1005-treated female mice, indicating suppression of both canonical and non-canonical NF-κB signaling pathways (**Fig. 5D**). Further, the histopathological examination of lung tissues from female mice revealed findings consistent with molecular observations and closely aligned with those observed in male counterparts. Uninfected controls exhibited intact alveolar architecture with no evidence of inflammation. In contrast, vehicle-treated SARS-CoV-2-infected female mice showed extensive alveolar disruption, dense inflammatory infiltration, and compromised pulmonary structure. Treatment with VLX-1005 substantially ameliorated these lesions, preserving alveolar integrity and markedly reducing inflammatory cell accumulation (**Fig. 5E**). These findings reinforce the role of 12-LOX in driving pulmonary inflammation during SARS-CoV-2 infection and demonstrate that its inhibition modulates distinct immune and inflammatory gene networks in a sex-dependent manner.

**Fig 5.**
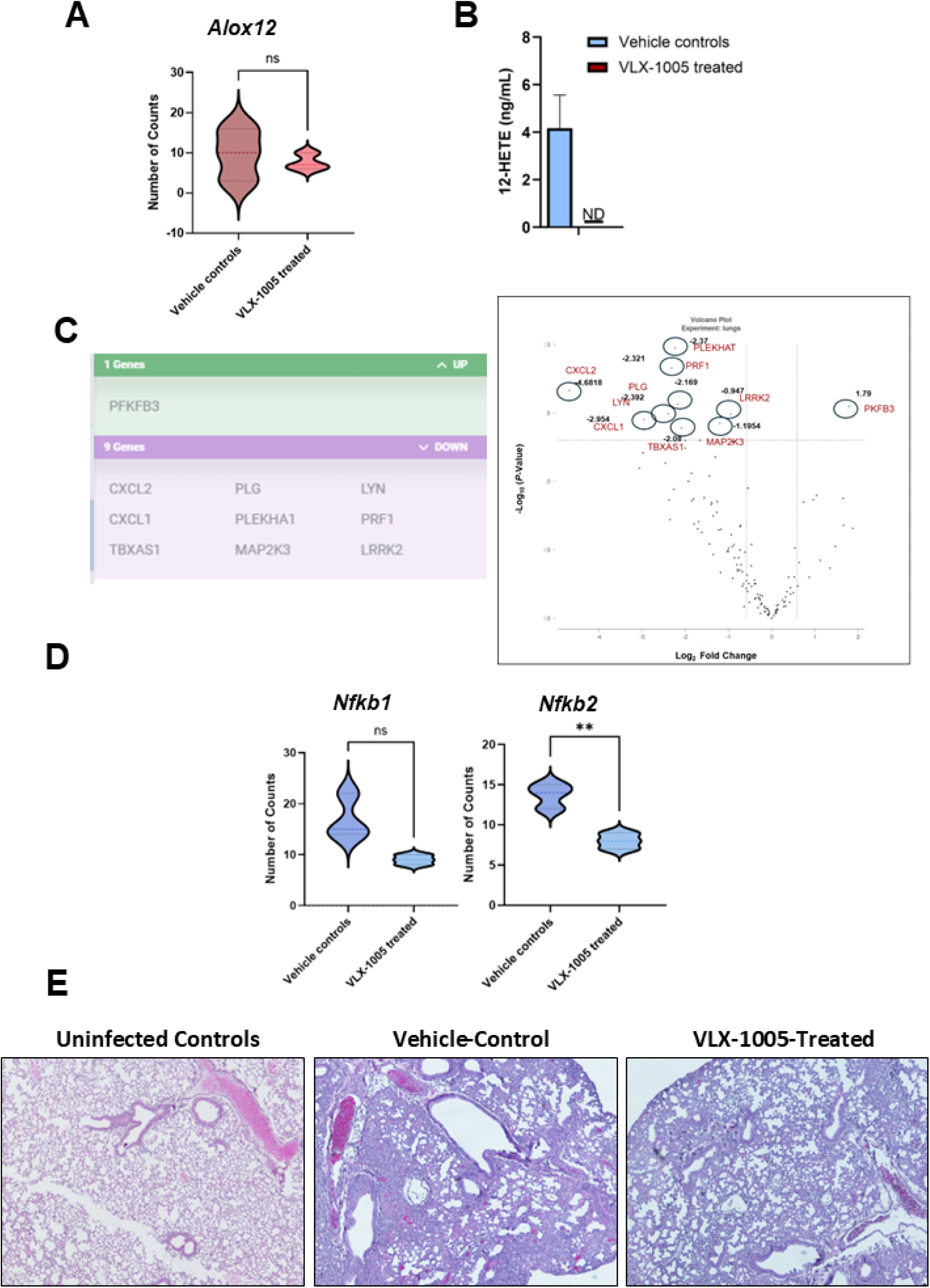
Inhibition of 12-LOX reduces inflammatory gene expression and lung pathology in SARS-CoV-2-infected female mice. Female K18-hACE2 transgenic mice were intranasally infected with 1 × 10^5^ PFU of SARS-CoV-2 and treated intraperitoneally with either vehicle or VLX-1005 (30 mg/kg/day) from 2 to 14 days post-infection. All VLX-1005-treated mice were euthanized, while vehicle-treated mice were necropsied immediately upon mortality for sample collection. Lung tissues and serum were harvested for molecular analyses. **(A)** NanoString quantification of *Alox12* mRNA levels in lungs (n = 3 mice per group). **(B)** 12-HETE levels in serum as determined by ELISA (n = 5 mice per group). **(C)** Summary of differentially expressed genes identified by NanoString analysis (n = 3 mice per group) shown in circles as Log₂ fold-change in expression. **(D)** Expression of *NfkB1* and *NfkB2* genes in lung tissues (n = 3 mice per group). The data were analyzed using Student’s *t-test.* *p<0.05. **(E)** Representative H&E-stained lung sections from uninfected controls, SARS-CoV-2-infected vehicle-treated, and VLX-1005-treated mice (n = 3 mice per group).

## Discussion

Previous studies have highlighted the central role of 12-LOX in orchestrating inflammatory responses in metabolic disease and suggest that targeting this pathway may offer dual benefits in modulating both immune and metabolic dysfunction. The enzyme 12-LOX, encoded by the *ALOX12* gene, plays a pivotal role in the pathogenesis of diabetes by promoting chronic inflammation through the generation of bioactive lipid mediators such as 12-HETE (24). In both type 1 and type 2 diabetes, 12-LOX expression is upregulated in pancreatic islets, adipose tissue, and immune cells, contributing to the inflammatory milieu that underlies metabolic dysfunction (25). Mechanistically, 12-LOX activity enhances the production of reactive oxygen species (ROS), promotes endoplasmic reticulum stress, and activates pro-inflammatory transcription factors such as NF-κB via 12-HETE, leading to increased secretion of pro-inflammatory cytokines (10, 26). In pancreatic β-cells, 12-HETE has been shown to impair insulin secretion and promote apoptosis, thereby exacerbating β-cell dysfunction and accelerating disease progression (27). Moreover, lipidomic analyses have revealed an imbalance between pro- and anti-inflammatory lipid mediators in diabetic tissues, with elevated levels of 12-LOX products contributing to the failure of resolution pathways (7). Importantly, pharmacological inhibition or genetic deletion of 12-LOX in preclinical models has been shown to reduce inflammation, preserve β-cell mass, and improve glucose tolerance, underscoring its potential as a therapeutic target in diabetes (8). In this study, we investigated whether SARS-CoV-2 infection upregulates 12-LOX activity, thereby contributing to hyperinflammation, tissue injury, and mortality, and evaluated the therapeutic efficacy of VLX-1005, a selective 12-LOX inhibitor, in improving disease outcomes in a preclinical model of COVID-19.

Our study reveals a significant upregulation of 12-LOX in lung tissues from diabetic individuals who succumbed to COVID-19, compared to diabetic individuals who died from unrelated respiratory illnesses. Immunofluorescence staining showed strong ALOX12 protein expression in alveolar and bronchial epithelial cells of COVID-19-positive lungs, while COVID-19-negative diabetic lungs exhibited minimal staining. These protein-level findings were corroborated by transcriptomic analysis, which demonstrated elevated *ALOX12* mRNA levels in COVID-19-positive diabetic lungs. These results suggest that SARS-CoV-2 infection, rather than diabetes alone, drives the activation of the 12-LOX pathway in the lung. In our study, the upregulation of *ALOX12* was accompanied by increased expression of immune and inflammatory genes, including *CXCR2*, *CCR7*, *IL6R*, *IFIT3*, *OASL*, and *STAT2*. This transcriptional profile suggests a coordinated inflammatory response involving chemokine signaling, cytokine receptor activation, and interferon-stimulated gene expression.

Our results demonstrate that SARS-CoV-2 infection alone is sufficient to induce pulmonary 12-LOX expression independent of metabolic comorbidities in the K18-hACE2 mice. Importantly, expression of 12-LOX, but not of other lipoxygenases such as 15-LOX, highlights the selective activation of the 12-LOX pathway in response to SARS-CoV-2 infection. Prior studies have shown that SARS-CoV-2 infection leads to mitochondrial dysfunction, oxidative stress, and activation of inflammatory signaling pathways in lung epithelial and endothelial cells, which are known to activate redox-sensitive transcription factors such as NF-κB, which in turn can upregulate *ALOX12* expression (28–30). These observations, integrated with our transcriptomic data from diabetic human lung tissues from deceased COVID-19-positive individuals, support a model in which SARS-CoV-2 acts as a potent inducer of 12-LOX-mediated inflammatory signaling.

A highly selective 12-LOX inhibitor, VLX-1005, previously known as ML355 (29), was discovered through a comprehensive screening of approximately 150,000 compounds and multiple rounds of iterative medicinal chemistry optimization. Pre-clinical safety testing with VLX-1005 exhibits excellent potency against 12-LOX (IC_50_ 340 nM) and displays remarkable selectivity against the related enzymes in the arachidonic acid pathways, including 15-LOX-1 (∼30 fold), 15-LOX-2 (>290 fold), 5-LOX (>290 fold), and COX1/2 (>40 fold). VLX-1005 has no mutagenic activity and exhibits stability in mouse and human blood for over 2 and 4 hours, respectively. Another crucial aspect of VLX-1005 is its ability to prevent excessive thrombosis, and Phase 2 clinical trials are underway to test the safety and tolerability of single and multiple intravenous doses of VLX-1005 for the treatment of heparin-induced thrombocytopenia.

This study provides compelling evidence that 12-LOX activity significantly contributes to the pathogenesis of SARS-CoV-2 infection in K18-hACE2 mice, and that pharmacological inhibition of this pathway using VLX-1005 confers a protective effect. The therapeutic benefit of VLX-1005 was evident in both male and female mice, although the magnitude of protection varied by sex and viral inoculum. At lower viral doses, female mice exhibited a more resistant phenotype, characterized by minimal morbidity and complete survival in the absence of treatment. In contrast, male mice demonstrated greater susceptibility, with VLX-1005 treatment resulting in improved survival and attenuated weight loss. These findings are consistent with clinical observations in human populations and our previously published study, where males are disproportionately affected by severe COVID-19 outcomes, as compared to females (32–35).

At the molecular level, VLX-1005 treatment in male mice led to a significant reduction in both *Alox12* mRNA expression and its enzymatic product, 12-HETE, confirming effective inhibition of 12-LOX at transcriptional and functional levels. This is particularly relevant given the established role of 12-HETE in promoting oxidative stress and pro-inflammatory signaling in viral and metabolic diseases. Transcriptomic profiling revealed that 12-LOX inhibition resulted in broad immunomodulatory effects, including downregulation of 30 inflammation-associated genes. These included chemokines (*Ccl11, Ccr7, Cxcl2, Cxcl5, Cxcr2*), cytokine receptors (*Il6ra, Il1rap, Il1r2*), and immune modulators (*Map3k8, Socs3, Tap1*), all of which are implicated in the cytokine storm associated with severe SARS-CoV-2 infection (36–40). Notably, suppression of *Nfkb1* and *Nfkb2* transcripts in VLX-1005-treated mice suggests attenuation of both canonical and non-canonical NF-κB signaling pathways. Given the central role of NF-κB in amplifying inflammatory responses, its downregulation supports the hypothesis that 12-LOX inhibition mitigates the transcriptional machinery driving cytokine overproduction. These findings align with previous reports identifying NF-κB activation as a key driver of lung inflammation and cytokine storm in COVID-19 (41). In addition to suppressing pro-inflammatory pathways, VLX-1005 treatment was associated with upregulation of genes involved in immune regulation and tissue repair, including *Fos*, *Klrd1*, and *Tgfβ*. This suggests that 12-LOX inhibition may not only limit inflammation but also promote resolution and recovery. Histopathological analysis corroborated these molecular findings, with lungs from VLX-1005-treated mice exhibiting preserved alveolar architecture and reduced immune cell infiltration compared to vehicle-treated controls.

Sex-specific differences in the transcriptional response to 12-LOX inhibition were also observed. While VLX-1005 effectively reduced 12-HETE levels in both sexes, the downstream gene expression profiles diverged. In male mice, inhibition of 12-LOX led to broad suppression of chemokines, cytokine receptors, and immune modulators, including *Ccl11, Cxcl2, Il6ra*, and *Socs3*. In contrast, female mice exhibited a more selective transcriptional response, with significant downregulation of *Cxcl1, Cxcl2, Lyn, Plg*, and *Map2k3*. Despite these differences, *Nfkb1* and *Nfkb2* were consistently downregulated in both sexes, indicating a shared mechanism of NF-κB pathway suppression. These sex-specific transcriptional signatures likely reflect underlying differences in immune regulation. Female mice are known to mount more balanced immune responses, characterized by enhanced antiviral activity and reduced pro-inflammatory cytokine production. This may explain the reduced disease severity observed in females at lower viral doses and the more targeted, yet effective, transcriptional response to VLX-1005. The upregulation of *Pfkfb3,* a bifunctional enzyme that regulates the key steps in glycolysis in female mice, further suggests a potential role for metabolic reprogramming in supporting immune resolution and tissue repair. Collectively, these findings highlight the critical role of 12-LOX in modulating the host inflammatory response to SARS-CoV-2 and support its therapeutic targeting as a strategy to mitigate disease severity. The observed sex differences underscore the importance of considering biological sex in the development and evaluation of host-directed therapies for COVID-19 and other viral infections.

In contrast to our findings, which demonstrate that 12-LOX contributes to inflammation, germline deletion of platelet *Alox12* has been reported to worsen disease outcomes, implying a protective role for 12-LOX-derived lipid mediators in the context of SARS-CoV-2 infection (42). Compensatory mechanisms may explain this paradox: deletion of *Alox12* leads to upregulation of *Alox15*, increasing 12/15-LOX activity and amplifying the production of inflammatory lipid mediators such as 12-HETE and 15-HETE. The resulting increase in 12/15-LOX activity therefore exacerbates inflammation and worsens disease severity in *Alox12*-deficient mice (43, 44). In contrast, as observed in this study, VLX-1005 selectively inhibits 12-LOX enzymatic activity without altering gene expression, thereby avoiding the compensatory upregulation of 12/15-LOX observed in genetic knockout models. This pharmacological approach offers a more precise and controlled strategy for modulating inflammation, particularly in diseases like COVID-19, where immune dysregulation contributes to pathology.

In summary, this study identifies 12-LOX as a key regulator of pulmonary inflammation in SARS-CoV-2 infection and validates its inhibition via VLX-1005 as a viable therapeutic strategy. VLX-1005 not only attenuated the cytokine storm by reducing pro-inflammatory mediators but also preserved lung architecture, indicating dual anti-inflammatory and tissue-protective effects. These findings support further investigation into 12-LOX inhibition as a host-directed approach for managing COVID-19-associated lung injury and potentially other inflammatory respiratory diseases.

## Acknowledgements

We gratefully acknowledge Dr. Preeti Bharaj, Dr. David Alland, and the staff at the Rutgers Regional Biocontainment Laboratory for their expert assistance in conducting the in vivo mouse experiments. We also thank Dr. Anjali Saki at Columbia University BioBank for their support in providing human autopsy specimens. This work was funded by a research grant from Veralox Therapeutics Inc.; the sponsor had no involvement in the study design, data acquisition, analysis, interpretation, or manuscript preparation.

